# A novel lncRNA PROCA11 regulates cell plasticity in response to androgen deprivation of prostate cancer cells

**DOI:** 10.1101/2023.08.07.552355

**Authors:** Rocco Cipolla, Marc Gabriel, Giorgia Ianese Regin, Micaela Piemontese, Ugo Szachnowski, Virginie Firlej, Marina Pinskaya, Antonin Morillon

**Author notes:** co-corresponding and co-last authors.

## Abstract

Long non-coding RNAs (lncRNAs) represent vast and yet poorly characterized family of genes that can fine tune cellular plasticity, thereby allowing the emergence of aggressive therapy-resistant and metastatic cancers. Androgen deprivation therapies (ADT) are commonly used to treat prostate cancer by inactivating the Androgen Receptor (AR). However, castration-resistant prostate cancer (CRPC) with neuroendocrine subtypes (NEPC) often emerge. In this study, we explore the role of lncRNAs in response to androgen deprivation. Using a dynamic prostate cancer cell system mimicking the CRPC and NEPC onset, we identified 15 novel lncRNAs, with PROCA11 standing out as a first-choice candidate, being also highly abundant in high-risk prostate cancer tumors. This majorly nuclear lncRNA is expressed at low levels in androgen-dependent conditions of growth and strongly activated upon hormone withdrawal, preceding neuroendocrine genes and persisting at high levels in neuroendocrine cells. Extensive computational analysis of clinical data and functional studies in cells revealed PROCA11 association with basal-to-luminal transformation of the transcriptomic landscape and activation of metabolic and signaling pathways reminiscent of neurogenesis and of maintenance of AR signaling. We propose that PROCA11 is involved in the intricate circuits regulating cellular plasticity enabling cell survival and proliferation and emergence of the NE phenotype in response to ADT.

## Introduction

Initiation and progression of prostate cancer depend on androgens and on the androgen-inducible transcription factor - Androgen Receptor (AR). Hence, androgen deprivation therapies (ADT) in combination with AR antagonists are used as a first-line treatment of prostate cancer. However, patients often relapse and develop castration-resistant prostate cancer (CRPC), dropping the 5-years survival rate to 30% (Siegel *et al*, 2023). Furthermore, up to 20% of CRPC patients undergo histological transformation from adenocarcinoma to neuroendocrine subtype, also known as therapy-induced neuroendocrine prostate cancer (tNEPC), which is considered lethal (Beltran *et al*, 2011); (Abida *et al*, 2019); (Yamada & Beltran, 2021). CRPC is characterized by high heterogeneity in tissue morphology, cellular and molecular characteristics. Emerging single-cell and bulk transcriptomic studies of post-ADT tumors reveal high heterogeneity with a complex combination of cell types for majority retaining properties of prostate adenocarcinoma with still active AR signaling, but some progressively acquiring androgen independence (AI) and/or undergoing neuroendocrine (NE) trans-differentiation giving rise to tNEPC (Patel *et al*, 2019) ; (Beltran *et al*, 2019) ; (Beltran *et al*, 2012) ; (Labrecque *et al*, 2019) ; (Risbridger *et al*, 2021). However, these trajectories are not linear and prevailing, reflecting high lineage plasticity of advanced prostate cancer tumors due to multiple genetic alterations and activation of numerous reprogramming transcription factors (ONECUT2, BRN2, SOX2, OCT4, ASCL1) and chromatin modifies (EZH2, SWI/SNF) in response to inhibition of the AR signaling (Choi *et al*, 2022); (Han *et al*, 2022); (Nouruzi *et al*, 2022); (Bishop *et al*, 2017) ; (Cyrta *et al*, 2020).

Recently massive RNA-sequencing revealed thousands of long noncoding (lnc)RNAs, constituting a major family of genes in humans, highly expressed in CRPC and some in NEPC tumors (Abida *et al*, 2019); (Ramnarine *et al*, 2018) ; (Iyer *et al*, 2015). Among them, MIAT, lncRNA-p21, LINC00261 and H19 were described as regulators of the NE trans-differentiation in prostate cancer (Crea *et al*, 2014); (Luo *et al*, 2019); (Singh *et al*, 2021); (Mather *et al*, 2021). These lncRNAs were shown to contribute to tumor plasticity and cancer progression modulating gene expression patterns mostly at the transcriptional level. MIAT, a myocardial infarction associated transcript, and H19 are up-regulated in NEPC and proposed to interact with and regulate the activity of PRC2 reshaping H3K27me and H3K4me3 at gene promoters towards neuroendocrine trans-differentiation (Crea *et al*, 2014); (Singh *et al*, 2021). Another study revealed that the treatment of CRPC patents with the AR antagonist enzalutamide induces the expression of lncRNA-p21 enabling switching the EZH2 methyltransferase activity from the histone H3K27 to a non-histone substrate, the STAT3 transcription factor (Luo *et al*, 2019). Finally, LINC00261 is an example of an evolutionary conserved lncRNA, which contributes to proliferation and metastasis in NEPC through two distinct nuclear and cytoplasmic mechanisms (Mather *et al*, 2021). Apart from these unique examples, our knowledge of how lncRNAs enable to circumvent the chemical castration in the onset of CRPC and NEPC stays very limited.

In this study we took advantage of the dynamic prostate cancer cell system recapitulating the CRPC onset to identify lncRNAs induced early in response to androgen deprivation. Total RNA-sequencing followed by differential expression analysis of *scallop*-assembled transcripts and multiple filtering isolated 15 novel lncRNAs, including the top candidate PROCA11, which are highly expressed in high-risk prostate cancer tumors and associated with emergence of NEPC traits. This majorly nuclear lncRNA, presenting 3 main isoforms, is lowly expressed in hormone-dependent cells but activated following AR knock-down or due to a release of AR from *PROCA11* promoter upon androgen deprivation. Activation of PROCA11 transcription precedes those of NE genes and persists at high levels though NE trans-differentiation. Gain- and loss-of-function experiments demonstrate that PROCA11 promotes cell proliferation regulating a balance between cAMP and RAS signaling, synaptic and metabolic activities. We propose that in the context of androgen deprivation of prostate cancer cells, PROCA11 can modulate metabolic profile and potentiate cells for survival and proliferation in the route of acquisition of neuroendocrine phenotype.

## Results

### Discovery of a novel lncRNA PROCA11, induced by androgen deprivation of prostate cancer cells

To characterize transcriptome changes underlying cell plasticity in response to androgen deprivation we took advantage of a dynamic *in vitro* cell model recapitulating early response to ADT androgen deprivation and progressive acquisition of NE-like and AI traits (Terry *et al*, 2013). In this system, first, adenocarcinoma LNCaP cells are grown in the presence of dihydrotestosterone (DHT) mimicking primary epithelial androgen-dependent luminal tumors (AD-cells). Then, they are subjected to androgen deprivation which induces immediate growth arrest, progressive change in cell morphology and properties towards the neuroendocrine-like phenotype (NE-like cells) with weaker but still persisting AR in the nucleus (Figure 1A; Supplementary Figure 1A-B). We performed a total stranded RNA-sequencing of these cell populations at different time points of transition and quantified gene expression using a custom transcriptome annotation including the Gencode release 32 and *scallop*-assembled transcripts from intergenic and antisense regions and regions overlapping annotated noncoding sense exonic transcripts (see Material and Methods). To focus on the earliest changes induced by androgen deprivation we further performed a differential expression analysis comparing two conditions: AD-cells actively proliferating in the presence of DHT and growth arrested NE-like cells cultured for 15 days and 1 month in hormone-depleted media (NE-15d/1m) (Figure 1A). In total, 964 lncRNAs were significantly induced by androgen deprivation (adj. p-value ≤ 0.05, Figure 1B). Further selection of the most robust lncRNA candidates was done through a three-fold filtering: normalized read counts above 95^th^ percentile in NE-15d together with (ii) a significant increase of expression in high-risk versus low-risk prostate tumors of prostatectomy origin (adj. p-value ≤ 0.05) (Pinskaya *et al*, 2019); and (iii) co-expression with protein-coding genes functionally linked to neurogenesis (Pearson’s *r* ≥ 0.99) (Figure 1B). This allowed isolation of four already annotated and fifteen novel lncRNAs with the top most up-regulated candidate, named PROstate CAncer lncRNA 11 (PROCA11) (Figure 1C).

**Figure 1.**
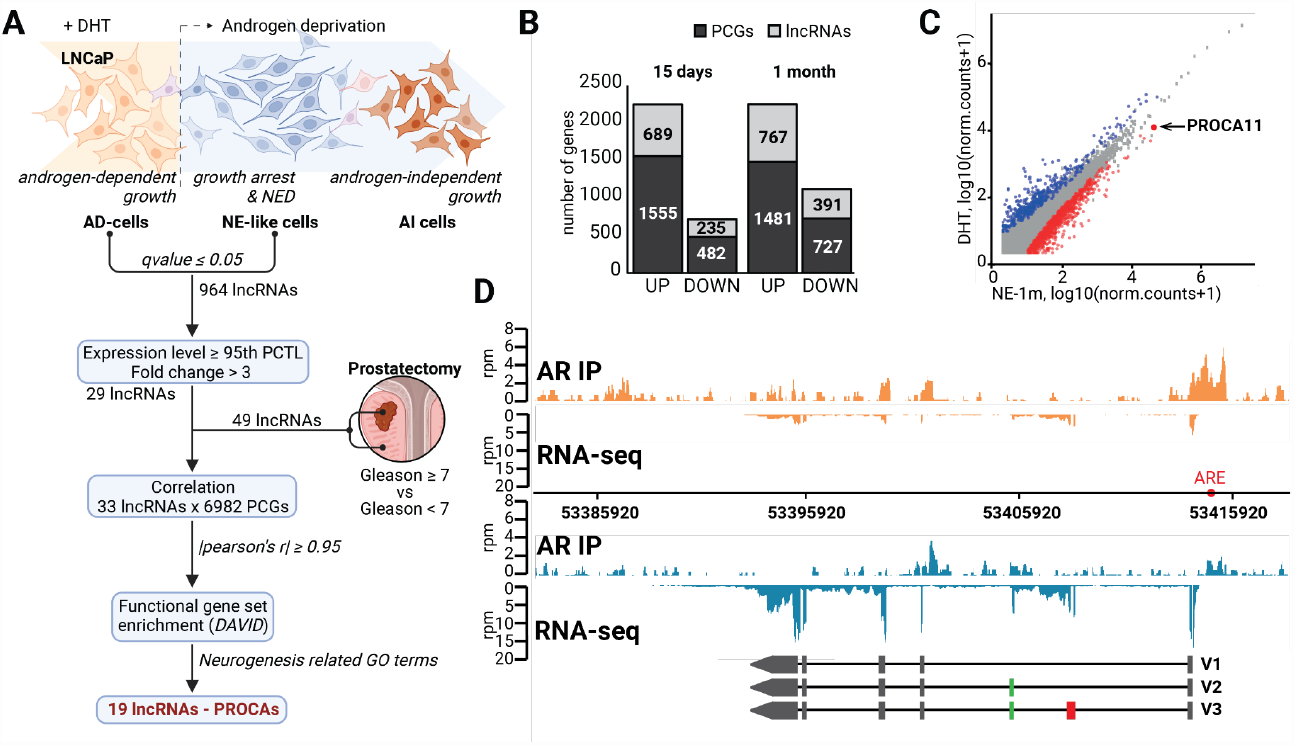
Discovery and characterization of novel PRostate Cancer Associated (PROCA) lncRNAs, expressed in response to androgen deprivation: (A) Experimental and bioinformatics workflow for discovery of lncRNAs associated with the onset of the NE phenotype (created with BioRender.com); (B) Significantly deregulated protein-coding (PCGs) and lncRNA genes identified in NE-like cells after 15 days and 1 month of androgen deprivation (adj. p-value ≤ 0.05); (C) 2D-plot of gene expression in log10(normalized counts) showing down-regulated genes (blue) and up-regulated genes (red), including PROCA11, upon 1 month of androgen deprivation; (D) PROCA11 RNA-seq and AR CUT&RUN profiling in AD-(orange) and NE-1m (bleu) cells across the chromosome 8 region embedding a newly annotated gene, genomic position of the putative ARE (GGACAAGGA) is indicated as a red dot. PROCA11 isoforms V1, V2, and V3 identified by molecular cloning and Sanger sequencing are represented with exons shown as rectangles, and introns as lines.

**Supplementary Figure 1.**
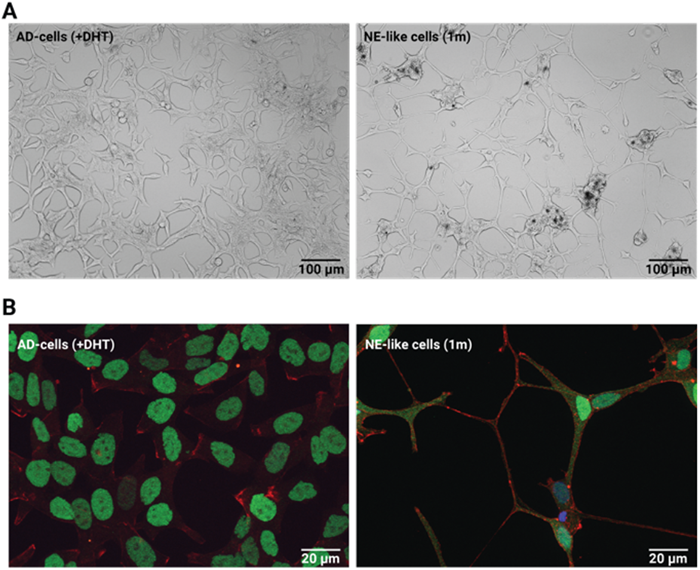
Androgen deprivation induces changes in cell morphology and AR subcellular localization: (A) Bright-field inverted microscope images of LNCaP cells growing in the presence of DHT (AD-cells) and in the absence of hormone for one month (NE-like cells, NE-1m); (B) Immunofluorescence imaging of AR (green), F-actin stained by phalloidin (red) and DNA stained by DAPI (bleu) in AD- and NE-1m cells.

Molecular cloning and sequencing identified three major isoforms of PROCA11, differing in inclusion of the second and third exons, that we named PROCA11 V1, V2 and V3, respectively (Figure 1E). Since PROCA11 expression was low in AD-cells but strongly stimulated in NE-like cells, we hypothesized that its transcription can be regulated by AR. Indeed, CUT&RUN profiling revealed a presence of AR at the *PROCA11* promoter, enclosing an *in silico* predicted androgen response element (ARE) (Messeguer *et al*, 2002), in AD-cells growing in the presence of androgen. The AR peak was dramatically decreased upon androgen deprivation concomitantly with the increase in PROCA11 levels (Figure 1D).

To understand the relationship between AR and PROCA11 expression we performed RNAi-mediated depletion of AR in AD-cells. It resulted in a 30% increase of PROCA11 levels in contrast to the prostate specific antigen (PSA)-encoding *KLK3* gene, a known AR-activated target (Figure 2A). We, next, isolated nascent transcripts from the chromatin fraction of cells treated with alpha-amanitin to block transcription. Further quantification of intronic sequences revealed a substantial increase of PROCA11 nascent transcript levels and, hence, of its transcription upon androgen deprivation (Figure 2B). This response was very rapid, even preceding the detection of the characteristic NE marker - ENO2 (Figure 2C). Finally, we assessed a subcellular localization of PROCA11 by single-molecule RNA-FISH (smiFISH). In cells growing in the presence of the DHT hormone we detected in average 15 molecules of PROCA11 per cell increasing up to 2.5-fold upon hormone withdrawal for 72 hours (Figure 2D-E). In both conditions up to 80% of PROCA11 transcripts was localized in the nucleus (Figure 2F). Albeit, subcellular fractionation and quantification of specific PROCA11 isoforms revealed their uneven distribution with V1 and V2 being more abundant in the cytoplasm and V3 in the chromatin (Figure 2G).

**Figure 2.**
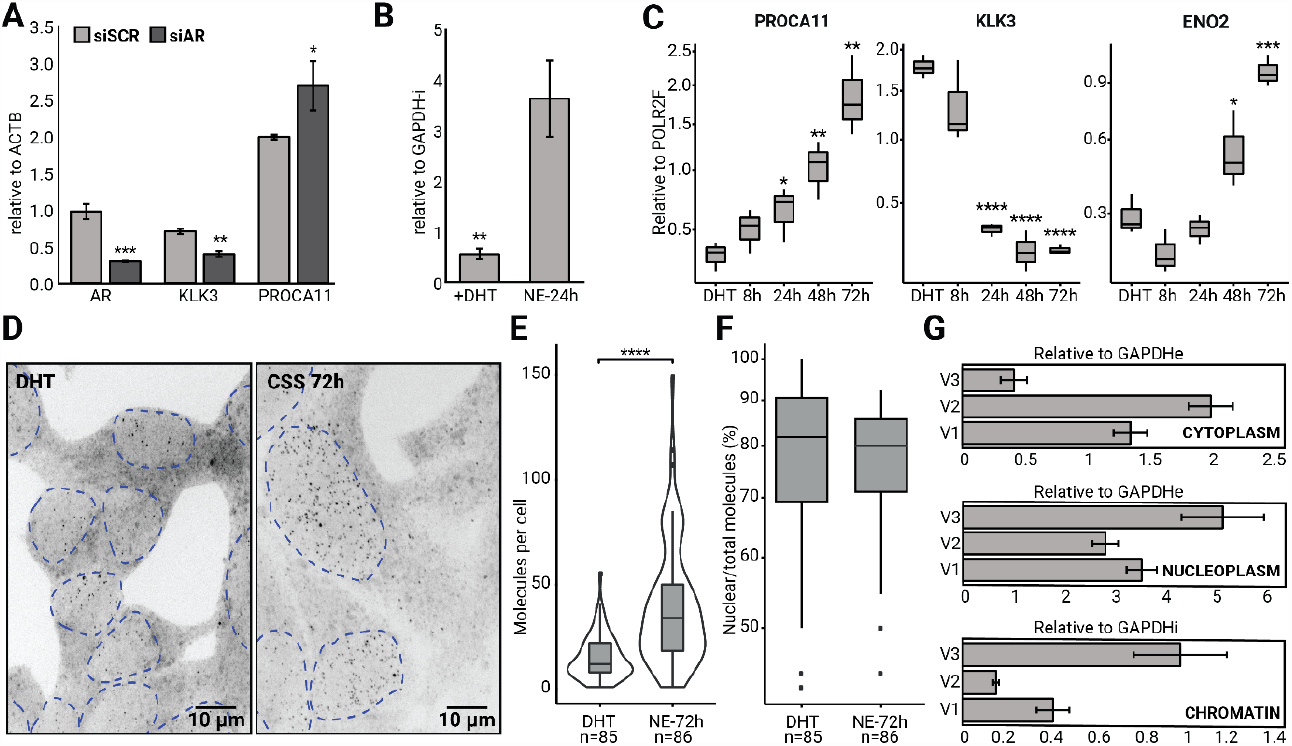
PROCA11 is an AR-repressed lncRNA transcriptionally activated very early upon androgen deprivation and localized majorly in the nucleus: (A) RTqPCR quantification of PROCA11 lncRNA, AR, and KLK3 levels in cells treated with siAR or siSCR 24 hours post-transfection, relative to siSCR; (B) RTqPCR quantification of nascent PROCA11 as intron relative to GAPDH intron in the chromatin fraction of alpha-amanitin treated cells growing in the presence or absence of DHT for 72h; (C) RTqPCR quantification of PROCA11, KLK3, and of the NE marker ENO2 in AD-cells in the presence of androgen and after 8, 24, 48, 72 hours of androgen deprivation, relative to POLR2F; (D) Representative smiFISH images of PROCA11 in cells growing in the presence and absence of hormone (DHT and NE-72h, respectively), dashed lines represent the outline of the nucleus; (E) Violin plot representation of the number of PROCA11 transcripts per cell in the DHT (N=85) and NE-72h conditions (N=86) (p-value < 4.5×10^−11^, Student t.test), and (F) Percentage of the PROCA11 nuclear foci per cell quantified by smiFISH and ImageJ. Asterisks indicate p-values range from 0.05 (*) to 0.0001 and below (****); (G) RTqPCR quantification of PROCA11 in cytoplasm, nucleoplasm and chromatin fractions of cells growing in the absence of androgen for 72 hours.

In conclusion, androgen deprivation of adenocarcinoma cells induces drastic transcriptomic changes with the most significant increase in levels of a novel lncRNA PROCA11. The *PROCA11* gene is composed of 7 exons that can be spliced in three variants, all showing mostly nuclear residence but distinct distribution between the nucleus and the cytoplasm. This can determine different fates and activities of PROCA11 isoforms in cells. Androgen deprivation leads to a release of AR from the *PROCA11* promoter and burst of its transcription very early in the onset of the NE phenotype. Therefore, we speculated that PROCA11 may contribute to acquisition of castration resistance as part of CRPC and NEPC transcriptomes.

### PROCA11 expression is elevated in ADT-treated adenocarcinoma, CRPC and NEPC tumors

To get insights into clinical relevance of the newly discovered lncRNA, we explored the expression of PROCA11 in numerous publicly available clinical datasets and compared it to our LNCaP system. First, the Genotype-Tissue Expression (GTEx) query revealed rather low PROCA11 expression in normal tissues (more than 100 times less) than in our primary adenocarcinoma cells with an exception of the ovary where it reached basal expression of the AD-cells growing in the presence of androgen (Carithers *et al*, 2015) (Figure 3A-B).

**Figure 3.**
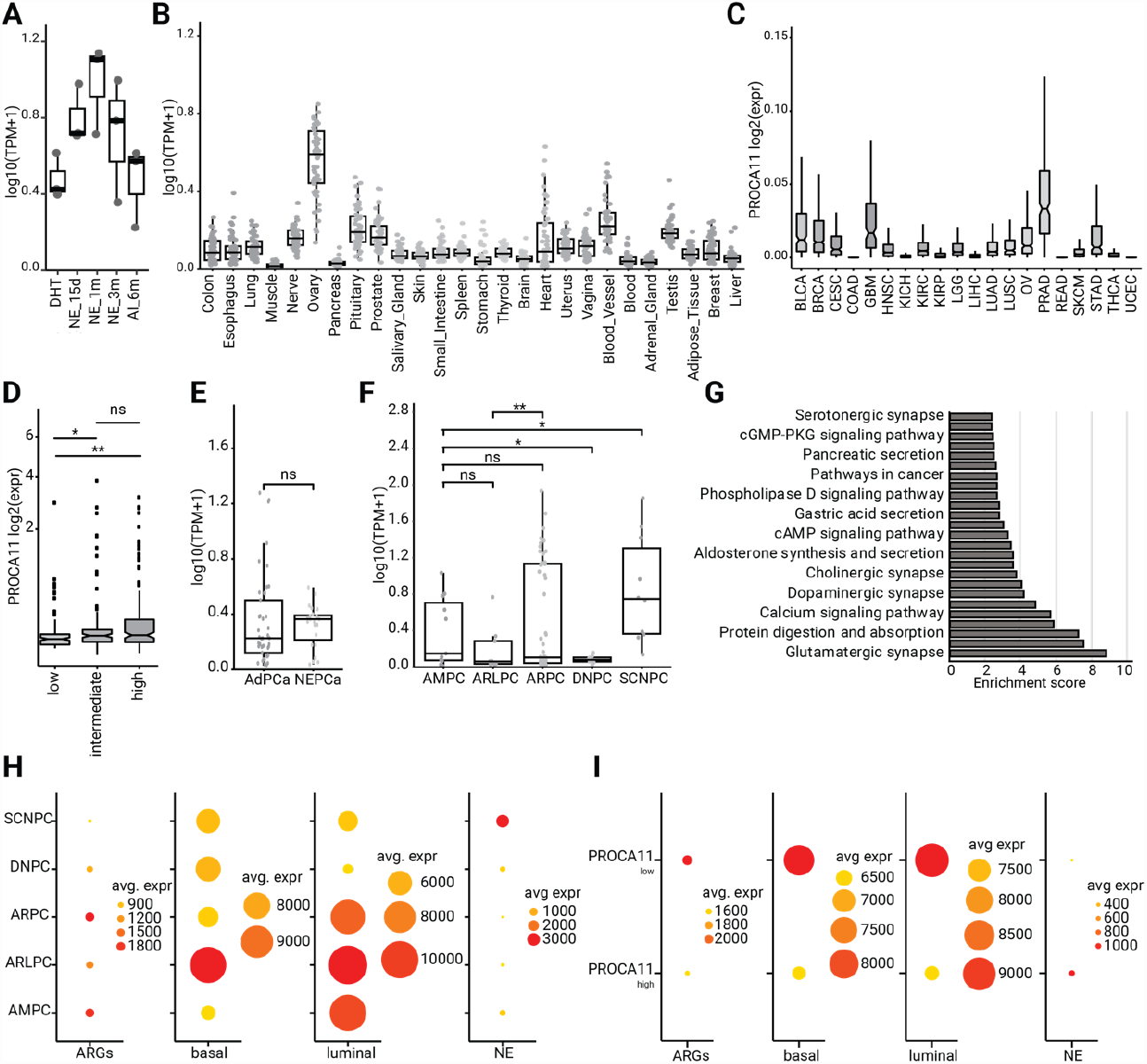
PROCA11 expression patterns in normal and tumor tissues suggest strong association with CRPC and NEPC onset: PROCA11 expression quantification in the dynamic LNCaP system of androgen deprivation (A), and across the GTEx collection of normal tissues (B) ; TCGA cohorts (C); TCGA-PRAD depending on tumor risk prognosis (D); AdPC and NEPC tumors from (Beltran *et al*, 2019) (E); and, finally, CRPC tumors from (Labrecque *et al*, 2019) (F); (G) KEGG pathways associated with up-regulated genes in tumors of AMPC, ARPC and SCNPC subtypes with high PROCA11 levels; (H-I) Dot plot of mean expression of genes constituting AR-positive targets - ARGs (Zhang *et al*, 2018), basal, luminal and NE signatures, retrieved from (Henry *et al*, 2018), in CRPC tumors from (Labrecque *et al*, 2019) and only AMPC and ARPC subtypes with high and low PROCA11 levels.

We next evaluated PROCA11 levels in cancer tissues across The Cancer Genome Atlas (TCGA) cohorts. As expected, it was markedly elevated in prostate adenocarcinoma (PRAD), but also notably high in glioblastoma multiforme (GBM), and moderately abundant in bladder urothelial carcinoma (BLCA) and breast invasive carcinoma (BRCA) (Figure 3C). Significantly, the higher lncRNA abundance in intermediate- and high-risk prognosis specimens of TCGA-PRAD than in low-risk tumors reinforced our hypothesis of PROCA11 association with advanced malignancy (Figure 3D).

Finally, PROCA11 expression was interrogated in metastatic CRPC with 41 tumors classified as castration-resistant adenocarcinoma with still active AR signaling (AdPC) and 23 tumors of neuroendocrine prostate cancer (NEPC) with inactive AR (Beltran *et al*, 2019), (Beltran *et al*, 2011). Remarkably, PROCA11 abundance was comparable in both subgroups (p-value < 0.05) (Figure 3E). We decided to explore more deeply a relationship between PROCA11, AR and NE genes signatures in another clinical study of post-ADT metastatic CRPCs subclassified into tumors with high expression of AR (ARPC, AR^+^/NE^-^), with low expression of AR (ARLPC, AR^low^/NE^-^), of small cell or neuroendocrine histology with expressed NE genes but lacking AR (SCNPC, AR^-^/NE^+^), double positive amphicrine tumors (AMPC, AR^+^/NE^+^) and, finally, double negative tumors lacking both AR and NE genes expression (DNPC, AR^-^/NE^-^) (Labrecque *et al*, 2019). Remarkably, PROCA11 expression was highest in SCNPC, but still detectable in all tumor subtypes, though surprisingly low in ARLPC and DNPC. This observation confirmed that PROCA11 is expressed in AR-positive cells, but also requires other factors or stimuli provided by NE cells. Although, it remains difficult to conclude if elevated PROCA11 expression is a consequence or a cause of the onset of the NE phenotype. Moreover, our attention was drawn to the observation of a bimodal distribution of the lncRNA in ARPC and AMPC tumors with still active AR signaling reaching the mean expression levels in AR-negative SCNPC (Figure 3F). Hence, we undertook a naïve search for a transcriptomic pattern discriminating the best PROCA11 high (above 75^th^ percentile) and low (below 25^th^ percentile) tumors of ARPC and AMPC subtypes together with the SCNPC tumors highly expressing the lncRNA (Supplementary Figure 2). Differential expression (FC above 2, p-value below 0.05) and GO term (FDR below 0.05) analyses across these specimens revealed, among others, up-regulation of genes, whose activities can be associated with proliferation and neurogenesis (cAMP and calcium signaling pathways, synapse) and with secretory pathways typical of luminal cells (Figure 3G). Furthermore, in addition to PROCA11 the top three protein-coding genes most discriminative between high and low PROCA11 tumors were neuron-specific *CNTNAP5* and *NRXN3*, both part of the neurexin family of genes with functions in cell adhesion and intercellular communication in the nervous system, and *MAP2* encoding a protein involved in microtubule stabilization during neurogenesis. Comparison of expression patterns for AR-targeted genes, basal, luminal and NE signatures clearly show that elevated PROCA11 levels are associated with higher expression of luminal genes in AR-positive cells and NE genes in SCNPC, and low PROCA11 levels are associated with higher basal component of AR-and NE-negative DNPC subtype (Figure 3H-I). Finally, AR-positive cells with low PROCA11 expression are characterized by mixed luminal-basal pattern (Figure 3I).

Together, these observations demonstrated that the high PROCA11 expression coincides with the expansion of the luminal signature and of the NE phenotype in CRPC tumors.

**Supplementary Figure 2.**
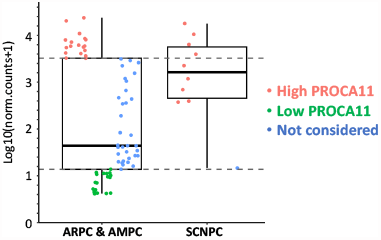
Assignment of PROCA11 expression levels: as high in SNPC (n=8), but also ARPC and AMPC subtypes (n=19) with lncRNA levels above 75^th^ percentile) (red) and as low in ARPC and AMPC subtypes (n=18, with lncRNA levels below 25^th^ percentile) (green). Specimens marked in red and green were used for differential expression analysis.

### PROCA11 can be expressed in adenocarcinoma and NE cells of an ADT-naïve tumor

PROCA11 expression in tumors with still functional AR signaling suggested that AR activity is not the only prerequisite governing lncRNA expression in CRPC tumors. High heterogeneity of tumors treated by ADT also led us to assume that the bulk RNA-sequencing misleadingly represents transcriptomes of distinct cell populations in the CRPC context. Hence, we decided to assess PROCA11 expression in two single-cell (sc)RNA-sequencing datasets from the MURAL collection of ADT-naïve patient-derived xenografts (PDXs) (Risbridger *et al*, 2021). One specimen recapitulated a mixed pathology with AR- and NE-positive cells (224R, AR^+^/NE^+^) and the other corresponded to a homogenous adenocarcinoma with active AR signaling and negative for NE genes (287R, AR^+^/NE^-^). A uniform manifold approximation and projection (UMAP) analysis separated 224R into two adenocarcinoma (AD) clusters and five NE clusters, and 287R into three adenocarcinoma clusters, comparably with the literature (Risbridger *et al*, 2021) (Figure 4A; Supplementary Figure 3B). Remarkably, PROCA11 was detectable only in the 224R mixed tumor, specifically, at high levels in one of the AD clusters, cluster 2, and parsimoniously in cells of all five NE clusters (Figure 4A). The 287R tumor showed low or any presence of the lncRNA (Supplementary Figure 3B). Hence, we concluded that PROCA11 is expressed in ADT-naïve tumors, as expected, in NE cells, but also in AD cells with still active AR signaling within a microenvironment created by NE cells. Further analysis of gene expression patterning across all clusters of the 224R tumor confirmed that PROCA11 is enriched in adenocarcinoma cells of more pronounced luminal cell type and lower basal signature, but also in cells with marked NE identity (Figure 4C). Gene ontology analysis of significant differentially expressed genes in the clusters 2 comparing to the cluster 3 revealed, in cells enriched in PROCA11 levels, notable down-regulation of translation (30 genes of RPL and RPS family, False Discovery Rate, FDR=6.45×10^−31^), and a tendency to down-regulation of the WNT signaling pathway (TIAM1, GPC3/6), pro-apoptotic, and cell cycle genes (*CDKN2A/p21, RACK1*). We speculated that AD-cells expressing PROCA11 have predominant luminal phenotype, enduring a particular low-proliferative state avoiding cell cycle arrest.

**Figure 4.**
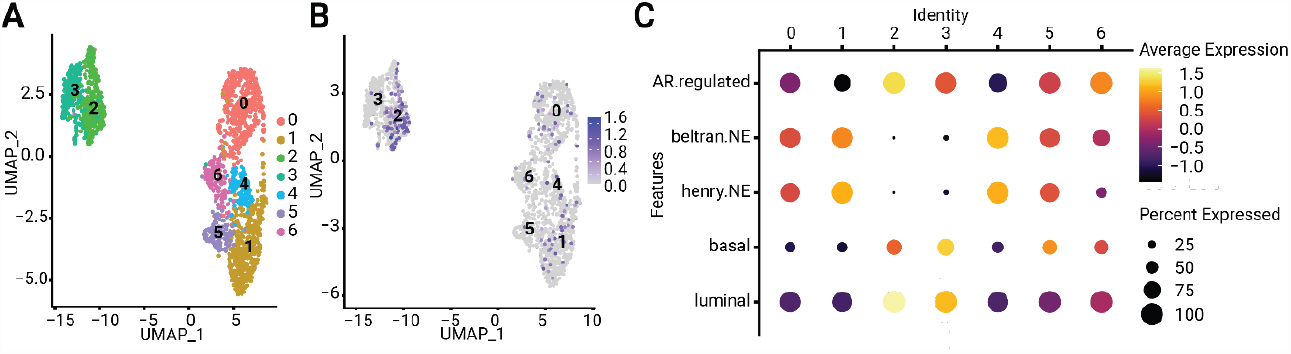
PROCA11 is expressed in the ADT-naïve tumor of the mixed adenocarcinoma and NE phenotype: UMAP (A - B) representation showing five NE and two AD clusters (A) and expression levels of PROCA11 (in log2(TP10K) based on scRNA sequencing (B); (C) Dot plot comparing expression of NE, basal and luminal gene signatures from (Henry *et al*, 2018) and (Beltran *et al*, 2016) across the AD and NE clusters for 224R.

**Supplementary Figure 3.**
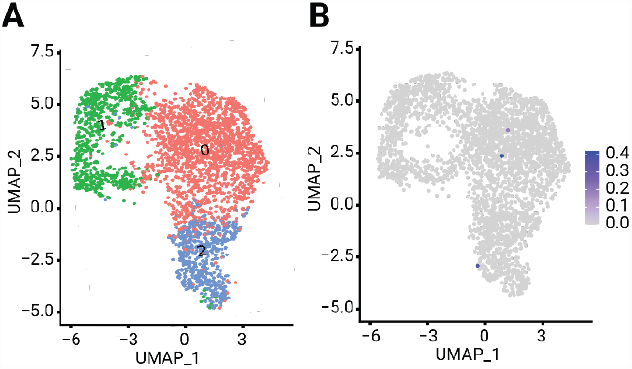
scRNA-seq analysis of the pure adenocarcinoma tumor revealed low or none PROCA11 expression: (A) UMAP of 287R separating cells into three AD clusters; (B) PROCA11 expression across 287R clusters quantified as log2(TP10K).

### PROCA11 regulates cell proliferation and cell cycle

Since in our model the increase of PROCA11 preceded neuroendocrine transdifferentiation and growth arrest, we hypothesized that it might have an effect on proliferation of cells. We thus performed gain- and loss-of-function studies to determine if alterations in lncRNA expression change cancer-related phenotypes. First, we sub-cloned the three major lncRNA variants into a lentiviral construct under the control of the constitutive CMV promoter and transduced them into LNCaP (oeP11) to overexpress each of three variants (oeV1, oeV2 and oeV3, respectively). To decrease PROCA11 levels we used an siPOOL strategy transfecting LNCaP cells growing in the presence or the absence of androgen with a mix of thirty siRNAs targeting exons common to all PROCA11 variants (siP11) or non-targeting controls (siSCR) (Figure 5A). PROCA11 depletion by siP11 was efficient achieving 80-90% of decrease in cells over-expressing PROCA11 variants though less robust (no more than 50%) in physiological conditions (Figure 5A). To follow cell proliferation, we used an xCELLigence real-time cell assay (RTCA) based on continuous measurements of electrical impedance. AD-cells and oeV1/V2/V3 cells overexpressing PROCA11 were seeded on plates, transfected or not with siRNAs and monitored during 260 hours for proliferation. Stable and constitutive PROCA11 overexpression resulted in faster proliferation rates for all three lncRNA isoforms (Figure 5B). And reverse, transient PROCA11 depletion in AD-cells and oeV1/V2/V3-cells reduced proliferation (Figure 5C).

**Figure 5.**
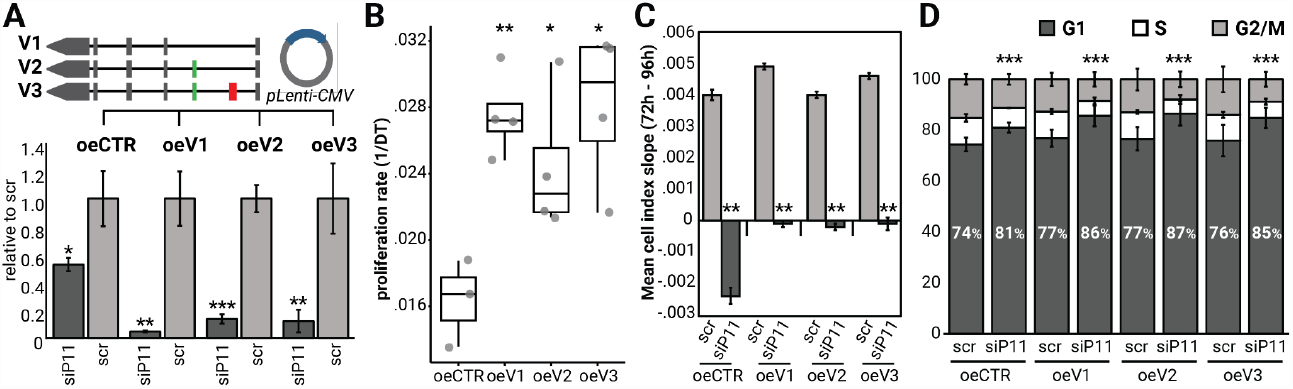
PROCA11 is involved in regulation of cell cycle and proliferation: (A) experimental setup for gain- and loss-of-function study of PROCA11 activity in AD-cells and RTqPCR quantification of PROCA11 levels in AD-cells and cells over-expressing PROCA11 (oeV1, oeV2, oeV3) transfected with siP11 relative to siSCR; (B) Proliferation rate (doubling time, DT^-1^) of oeCTR, oeV1, oeV2, oeV3 cells measured from 0 to 96 post-seeding ; (C) Cell index slope assessed for the exponentially growing cells transfected with siSCR or siP11 from 72 to 96 hours post-transfection; (D) Proportion of cells in G1, S and G2/M phases of the cell cycle. Asterisks represent the p-value range from 0.05 (*) to 0.001 and below (***) (Student t.test).

Proliferation depends on progression of a population of cells through the cell cycle. Hence, we assessed it in cells using flow cytometry and propidium iodide staining of DNA. PROCA11 depletion resulted in up to 10% increase in a proportion of cells in G1 over S and G2/M phases (Figure 5D).

Collectively, these results suggested that PROCA11 contributes to proliferation in the presence of androgen and may play a role in cell cycle progression. Although, all three isoforms seem to contribute similarly to lncRNA-induced phenotype changes.

### PROCA11 modulates metabolic and neurogenic activities in adenocarcinoma cells

In light of our findings that PROCA11 is expressed in AR-positive cells ubiquitously present in CRPC and that it regulates cell proliferation, we further investigated transcriptomic changes induced by the depletion of the lncRNA in AD-cells. Differential expression analysis of protein-coding genes identified 90 up- and 120 down-regulated genes (|FC| above 1.5, adj.p-value below 0.05). Strikingly, genes with increased expression were mostly part of Ras signaling, neurogenic activities, as glutamatergic and dopaminergic synapse, relaxin, cAMP and sphingolipid signaling pathways (Figure 6A). Notably, among up-regulated genes were several with important activities in neuron cell homeostasis and plasticity, as the MAPK10 signaling kinase, several genes encoding PLA2 enzymes of the fatty acid metabolism, or the DDC enzyme catalyzing the production of neural transmitters. On the other hand, down-regulated genes were associated with retinol metabolism and focal adhesion (Figure 6A). Among them, several genes belonged to the UGT2B family of UDP glucuronosyltransferases and to the cytochrome P450 family of enzymes, involved in the metabolism of different steroid hormones including androgens (Chouinard *et al*, 2007), (Mitra & Goodman, 2015). In addition, depletion of PROCA11 also led to an overall decline of the mean expression of the luminal cell-type signature and expansion of the basal signature (Figure 6B), as it was also determined by the scRNA-sequencing of the mixed AD/NE tumor in the AD cluster 3 with low or undetectable PROCA11 levels, but also in CRPC tumors with low PROCA11 (Figure 4C, Figure 3H-I).

**Figure 6.**
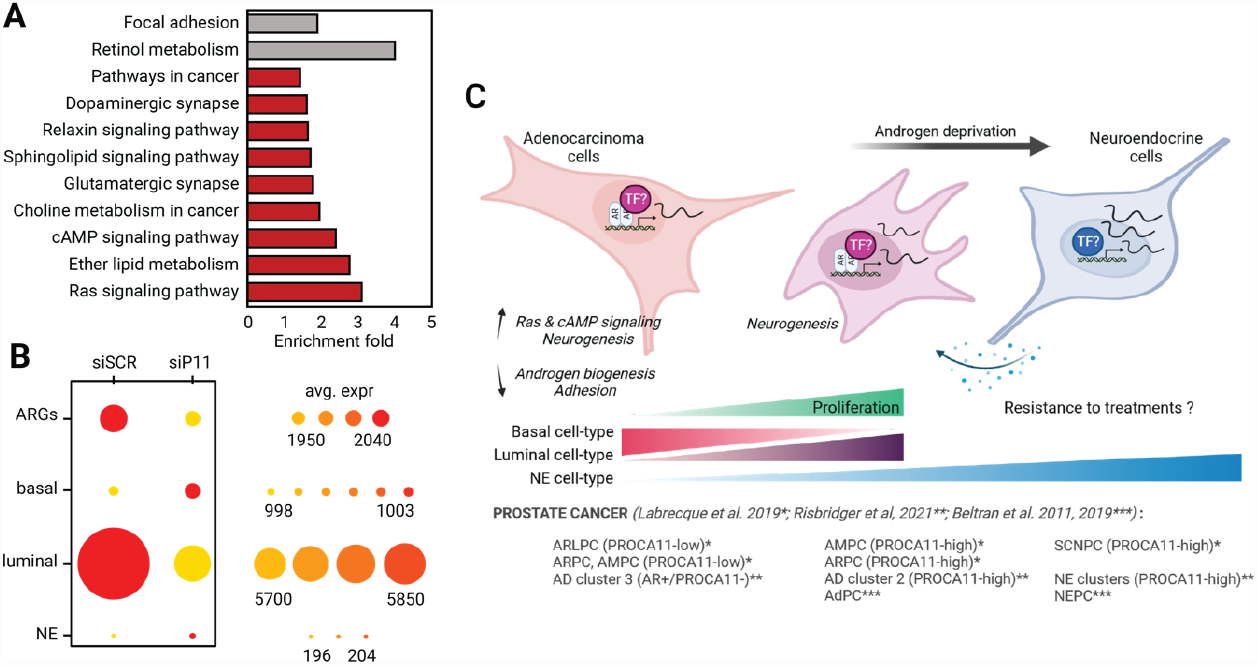
Alterations in PROCA11 expression change metabolism, adhesion and neurogenesis related pathways: (A) KEGG pathways enriched for up- and down-regulated genes (n=90 and 120, respectively) in response to PROCA11 depletion in AD-cells. Only genes with the expression |FC| above 1.5 were considered in GO term analysis by DAVID. (B) Dot plot comparing expression of NE and luminal gene signatures from (Henry *et al*, 2018) in AD-cells treated with siSCR or siP11; (C) Model of PROCA11 activity in the onset of CRPC and NEPC (created with BioRender.com).

In conclusion, our results strongly suggest that in cells growing in the presence of androgen PROCA11 contributes to the maintenance of the AR signaling to preserve a proliferative potential, modulates cellular plasticity switch from basal to more pronounced luminal cell type and potentiates cells to acquisition of NE properties.

## Discussion

Emerging examples suggest that lncRNAs intervene in fine tune regulation of many cellular processes modulating cell identity and fate (Chouinard *et al*, 2007), (Mattick *et al*, 2023). In cancer, lncRNAs are proposed to influence tumor progression and response to therapies (Carlevaro-Fita *et al*, 2020), (Vancura *et al*, 2021). In this work we discovered a novel lncRNA PROCA11 with a possible role in the onset of castration-resistant and neuroendocrine prostate cancer.

Transcriptome profiling of the dynamic *in vitro* system of the prostate adenocarcinoma revealed PROCA11 as the most activated in response to androgen deprivation. Importantly, PROCA11 levels are elevated in tumors of high Gleason grades, ADT-treated adenocarcinoma and NEPC tumors. Therapies targeting AR signaling, such as ADT, affect the plasticity of PCa cells by inhibiting the capacity of AR to activate transcription of proliferation and pro-survival genes. On the other hand, ADT results in a release of AR and activation of genes involved in stem-cell fate, epithelial-to-mesenchymal transition or neuroendocrine transdifferentiation (Beltran *et al*, 2016), (Nouri *et al*, 2017). Similarly, androgen ablation in the adenocarcinoma cell line induces, otherwise low, transcription of *PROCA11*. It coincides with the AR release from the lncRNA promoter and persists in growth arrested cells undergoing the neuroendocrine trans-differentiation. PROCA11 expression is also stimulated by the depletion of AR in the presence of androgen. Together, these findings strongly suggest either a negative regulation of lncRNA transcription by AR or through two distinct mechanisms: AR-mediated basal transcription in the presence of hormone, and strong activation by another transcription factor (TF) upon androgen withdrawal. *In silico* search for TF binding sites identified response elements of the basal TBP subunit of TFIID, of the AR, but also of the AP-1 complex members (JUN and FOS), several TFs (FOS, ESR1 and USF2), some other ligand-dependent TFs (GR, RAR/RXR), and differentiation TFs (POU2F2, LEF1, FOXA1), all of them are up-regulated in response to androgen deprivation. Interestingly, elevated ERɑ expression and downstream activation of its targets (TMPRSS2-ERG, PS2 or the NEAT1 lncRNA) were reported in AR-negative cells and PCa tumors (Setlur *et al*, 2008).

PROCA11 is not or very lowly abundant in almost all normal tissues with the exception of the ovary, but increases in advanced PCa tumors subjected to ADT, and specifically in tumors with NE features. ADT induces very heterogeneous changes in tumors, containing cells with still active AR signaling but also cells with inactive AR, cells acquiring stem-cell like or neuroendocrine features (Labrecque *et al*, 2019). In all studied clinical cases elevated PROCA11 levels were associated with ADT and co-expression of NE signature. However, scRNA-sequencing also revealed PROCA11 presence in the ADT-naïve tumor of the mixed adenocarcinoma and neuroendocrine phenotypes, but not in the pure homogeneous adenocarcinoma. Notably, it was expressed in AR-positive AD-cells presenting features of low or non-proliferative cells. In this context, PROCA11 may be part of a regulatory circuit preventing cell cycle arrest and apoptosis. One can also speculate that a particular NE microenvironment contributes to PROCA11 expression. NE cells secrete a variety of peptides and proteins with endocrine and paracrine effects, modulating the secretory activity of luminal cells to regulate prostatic growth and function (reviewed in Arman & Nelson, 2022). Therefore, in heterogeneous mixed tumors, NE cells’ secretome can activate a specific response, remodeling the transcriptome of adjacent adenocarcinoma cells stimulating PROCA11 expression and endowing them with the neuroendocrine properties and preserving their proliferative potential. Single-cell resolution studies in cells and tissues will shed light on the intricate relationship between PROCA11, the AR signaling and the onset of CRPC and NEPC phenotypes.

Depletion of PROCA11 in androgen-dependent cells leads to decreased cell proliferation and perturbed cell cycle with more cells in G0/G1 phase. Ectopic overexpression of PROCA11, in the contrast, promotes cell proliferation. Functional association analysis with GO analysis demonstrated that down-regulated genes are enriched for androgen biogenesis, as though up-regulated genes are involved in Ras and cAMP pathways. Activation of the Ras signaling in ADT treated prostate cancer was reported to reduce the androgen requirement of LNCaP cells for growth (Bakin *et al*, 2003). In a similar way, activation of the cAMP pathway was showed to promote AR translocation into the nucleus in conditions of low androgen levels, also stimulating AR signaling and survival (Dagar *et al*, 2019). Hence, activation of these two pathways in response to PROCA11 depletion converges to sensitize the AR activity to subphysiological levels of androgen. In the brain, cAMP function is tightly linked to synapse formation and neuron plasticity (reviewed in Shahoha *et al*, 2022). In PROCA11-depleted cells the cAMP pathway activation is concurrent with an increase in expression of genes involved in synaptic activities. These findings strongly point to the role of PROCA11 in regulation of cell plasticity enabling survival and growth but at the same time the emergence of the NE phenotype in conditions of perturbed androgen supply. This regulatory mechanism implies an intricate cross-talk between AR and PROCA11 and possible feedback loops with metabolic and neurogenic signaling pathways.

In the context of still functional AR signaling, elevated PROCA11 levels in our dynamic system, CRPC tumors and ADT-naïve tumor of the mixed AD/NE phenotype was associated with increased expression of luminal cell-type genes in detriment of basal cell-type genes. According to lineage tracing studies, basal cells have the ability to differentiate into luminal cells (Choi *et al*, 2012). Mixed basal-luminal (intermediate) population of cells resists to pharmacological castration and was suggested to given rise to CRPC. We speculate that in tumors PROCA11 may be involved in regulation of basal-luminal balance, contributing to the emergence of persister cells maintaining their proliferative potential and sensitized to neuroendocrine trans-differentiation. Therefore, ADT may induce the expression of PROCA11, first, as a compensatory response to counteract the growth arrest, and then enable to modulate cellular plasticity in favor of basal-to-luminal transformation and emergence of the NE phenotype.

The precise molecular mechanisms triggered by PROCA11 in the androgen-dependent and independent contexts still stay to decipher. So far, in *silico* predictions of the coding potential and studies of cell phenotypes induced by expression of different PROCA11 isoforms did not support their distinct fate and activity in cells. However, in light of recent findings on bifunctional lncRNAs, we cannot exclude that the shortest V1 and V2 isoforms, more enriched in the cytoplasm than V3, are translated, giving rise to a peptide with potential regulatory functions.

## Material and Methods

### Cell lines and culture

The LNCaP cell line was purchased from ATCC (CRL-1740), cultured at 37°C in a humid 5% CO2 atmosphere in either the RPMI 1640+GlutaMAX medium (Life Technologies) supplemented with 10% fetal bovine serum and 2 nM DHT (AD-cells, or +DHT) or the phenol red-free RPMI 1664+GlutaMAX medium supplemented with 10% dextran charcoal-stripped fetal bovine serum (NE- and AI-cells). HEK293T was purchased from ATCC (CRL-3216), cultures in DMEM + GlutaMAX (Life Technologies) supplemented with 10% fetal bovine serum. Cells were systematically tested and free of mycoplasma.

### Plasmids

The PROCA11 cDNA was obtained by reverse transcription and PCR with primers complementary to 5’ and 3’ extremities of the gene, V1, V2 and V3 amplicons were resolved in the 1% agarose gel and purified using the gel purification kit (Qiagen). Each cDNA was cloned into pDONR201 (Invitrogen) and further sub-cloned into pLenti CMV Blast DEST (706-1) (Addgene, #17451) (Campeau *et al*, 2009) using the Gateway system (Thermo Fisher Scientific). Constructs were validated by Sanger sequencing.

### Transfections and lentiviral transduction

Transfections were performed at 50-70% of cell confluency using the Jetprime® transfection kit (Polyplus) following the manufacturer’s protocol: (i) LNCAP with siPOOL (siTOOLs Biotech GmbH) targeting PROCA11 (siP11), AR (siAR) or non-targeting scramble (siSCR) at a final concentration of 30nM. Experiments were performed 48 hours post-transfection; (ii) HEK293T with 1.3 μg of psPAX2 (Addgene, #12260), 0.8 μg of pVSVG (Addgene, #36399) and 0.8 μg of the lentiviral plasmid hosting the cDNA of GFP lacking the first ATG (CTR) or PROCA11 variants (V1, V2, V3). 48 hours post-transfection, viral supernatant was collected, filtered and added to cells at 70% of confluency at 0.5 MOI for 24 hours. Transduced cells were subcultured in the medium supplemented with blasticidin at 6 μg/ml for four passages prior to use.

### Subcellular fractionation

The protocol was adapted from (Gagnon *et al*, 2014) for 10^7^ cells. All steps were performed on ice with ice-cold buffers supplemented with 20 U/µl SUPERase-IN (Thermo Fisher Scientific) for RNA extraction and 0.1 mM AEBSF (SIGMA) for protein extraction. For nascent transcription experiments, α-amanitin was added in the HLB buffer at 25μM concentration. RNA extraction from all fractions was performed using Qiazol at 65°C for 5 minutes with regular mixing and then purified with miRNeasy kit (Qiagen) according to the manufacturer’s instructions. For protein extractions, the chromatin fraction was sonicated for 4 min (30s-on/30s-off, “high power”) on the Bioruptor Plus (Diagenode). Proteins’ content was quantified with the Thermo Fisher Scientific− BCA Protein Assay Kit (Thermo Fisher Scientific).

### RNA extraction and reverse transcription

Total or AR-immunoprecipitated RNA extraction was performed directly from cell cultures with miRNeasy kit according to the manufacture’ instructions including the DNase treatment step (Qiagen). Reverse Transcription (RT) was performed on 500 ng of RNA with random primers and SuperScript II Reverse Transcriptase (Thermo Fisher Scientific).

### Quantitative PCR

cDNA was diluted 10-50 times and quantified by real-time PCR with LightCycler® 480 SYBR Green I Master (SIGMA). The *RPL11, POLR2F* or *ACTB* genes were used as reference genes to quantify a relative abundance of each cDNA. Error bars represent standard deviations in relative quantity of cDNA prepared from at least three independent experiments.

### Cell cycle analysis

In total, 200000 cells were recovered after trypsinization, washed twice in PBS, resuspended in 500*µ*l of ice-cold 70% EtOH and incubated at 4°C for 30 minutes. Subsequently, cell pellets were treated with 50*µ*l of RNase A solution (100*µ*g/mL) for 5 minutes at room temperature and stained in 300*µ*l of 50*µ*g/mL propidium iodide solution for the next 30 minutes. Cells were processed using the NovoCyte flow cytometer (Agilent). Data was acquired for 10000 single cells per condition and analyzed by the FlowJo software (v10.9.0).

### Proliferation

Cells were seeded (8000 cells/well) in a 96-well RTCA E-plate (Agilent) in 4 replicates per condition. Following a short spin at 130g for 1 minute, cells were transfected with siSCR or siP11 as described above. Electrical impedance was measured for 260 intervals of 1 hour using the RTCA xCELLIgence device (Agilent). Cell index slope and doubling time were retrieved from the software and used for statistical analysis in R.

### RNA-seq analysis

Read alignment files were used as input for running the Scallop v0.10.5 transcriptome assembler with default options (Shao & Kingsford, 2017). Assemblies were merged and mapped to the reference annotation Gencode Release 32 (GRCh38.p13) by the Cuffmerge v1.0.0 tool (Trapnell *et al*, 2010). All non-annotated transcripts with class code “u” (intergenic, n=4120 transcripts, n=2582 genes), “x” (antisense, n=2837 transcripts, n=1629 genes) and code “j” (sense exonic overlap, n=35 transcripts) restricted only to noncoding genes were retrieved and added to GRCh38.p13 to make a new custom transcriptome assembly. Reads were mapped using Tophat v2.0.4 (max mismatches allowed = 3) (Kim *et al*, 2013); (Trapnell *et al*, 2009). The uniquely mapped reads were counted at the gene level using featureCounts v2.0.0 (Liao *et al*, 2014) and the custom assembly build and, then, processed by DESeq2 v1.26.0 (Love *et al*, 2014) using the DESeq R package (Anders & Huber, 2010). Luminal, basal, NE gene signatures were retrieved from (Henry *et al*, 2018), ARGs were retrieved from (Zhang *et al*, 2018).

### Correlation expression matrix and gene ontology

A correlation matrix was built between each lncRNA and the totality of differentially expressed PCGs (n=6982) obtained from all the time points of hormonal deprivation (15 days, 1 month, 3 months, 6 months) using the pcor() in R. Highly correlated PCGs (|pearson r| >= 0.99) were selected for the gene ontology analysis with the R package clusterProfiler v3.14.3 (fun=enrichGO, ont=“BP”, pvalueCutoff =0.2). PROCA lncRNAs (n=15) make part of the functional group ([*nerv* | *neur* | *axon* | *synap*]).

### Public bulk and single-cell RNA-seq datasets query

Read counts from TCGA were retrieved by TANRIC (Li *et al*, 2015). Tumor classification as low-, intermediate or high-risk group was performed according to Gleason and TMN scores as previously described (Pinskaya *et al*, 2019).

Expression in normal tissues was obtained from the GTEx Portal via the SRA cloud data delivery service, by using the dbGaP eRA credentials for accession number phs000424.v9.p2, and the google cloud platform for storage on 12/15/2022 (https://gtexportal.org/home/) (Carithers *et al*, 2015). CRPC RNA-seq datasets were accessed through the NCBI GEO data repository under the number GSE126078 (Labrecque *et al*, 2019), and through dbGaP under the number phs001666.v1.p1 (Beltran *et al*, 2019). ScRNA-seq datasets of PDX 224R and 287R MURAL collection were downloaded from the Sequence Read Archive database (https://www.ncbi.nlm.nih.gov/sra/) under the BioProject accession PRJNA675382 and analyzed as described in (Risbridger *et al*, 2021).

### smiFISH

Probes design and procedure were carried out following the instructions of (Tsanov *et al*, 2016).

### Microscopy image analysis

Image analysis was carried using the ImageJ2 Fiji package (Schindelin *et al*, 2012). Stacks, spaced out by 0.6µm, were merged using *maximum intensity Z-projection*. Two regions of interest per cell defined total and nuclear areas. PROCA11 foci were counted using *find maxima* and the threshold = 80. Cytoplasmic counts were obtained by subtracting total with nuclear counts.

### CUT & RUN

The experiment was performed in two replicates following the protocol from Henikoff lab (Skene *et al*, 2018) with modifications described in (Jarroux *et al*, 2021). DNA fragments were purified with NEB Monarch PCR & DNA purification kit following the protocol enrichment for short DNA fragments and was quantified by Qubit HS DNA kit.

### CUT & RUN libraries preparation and sequencing

CUT & RUN libraries were prepared using the TruSeq ChIP-Sample Prep Kit-Set A and PCR Box (Illumina) from 5 ng of DNA with 10 PCR cycles for amplification following the manufacturer’s procedures. Concentration and quality were assessed by Qubit with dsDNA HS Assay Kit (Thermo Fisher Scientific) and High Sensitivity DNA Analysis Chip Kit (Agilent Technologies). Libraries were sequenced on a NovaSeq – S1 device in a pair-end 100 bp read length mode.

### CUT & RUN analysis

Reads were first aligned to human genome version GRCh38 using bowtie (version 2.4.1) (Langmead & Salzberg, 2012), with options -I 10 -X 700 --no-mixed. Reads with MAPQ value >=30 were kept for further analysis. To equilibrate datasets, the larger samples were randomly down-sampled. For AR, peaks were called relative to the IgG control using MACS2 version 2.2.6 (ref) and options: -f BAMPE -g hs -B --keep-dup all. For further analysis, common peaks between conditions were also merged to peak regions. The merged peaks were assigned to closest genes and the shortest distance between each peak and the closest transcription start site (TSS) coordinates were calculated using the Gencode v32 annotation. The AR binding site within the PROCA11 region was inferred using PROMO (Messeguer *et al*, 2002).

### Gene features analysis

Gene ontology (GO) biological process, KEGG pathways and cellular compartment terms enrichment analyses were performed using DAVID (Huang *et al*, 2008). The gene enrichment score was calculated as –log10(*P*-value).

## Acknowledgments

We acknowledge and thank Margot Hully, Dominika Foretek, Francis Vacherot and Damien Destouches for technical assistance and expertise. We thank the MURAL consortium and Dr Hamisha Beltran for their valuable datasets. This work was funded by ERC DARK and Malakof Humanis. R.C. was funded by the EuReCa PhD program and ARC.

## Author Contributions

Project administration, funding acquisition, conceptualization, cosupervision and writing, A.M.; conceptualization, cosupervision and writing, Ma.P.; draft revisions, Ma.P., R.C., A.M.; investigation R.C., Ma.P., V. F., Mi.P., G.I.R.; formal analysis, R.C., M.G., U.S.

**The authors declare no conflict of interest**.

